# Maxent modeling for predicting the potential distribution of global *talaromycosis*

**DOI:** 10.1101/2021.03.28.437430

**Authors:** Wudi Wei, Jinhao He, Chuanyi Ning, Bo Xu, Gang Wang, Jingzhen Lai, Junjun Jiang, Li Ye, Hao Liang

**Author notes:** indicate authors contribute equally (Wudi Wei, Jinhao He, Chuanyi Ning). indicate authors contribute equally (Li Ye, Hao Liang).

## Abstract

*Talaromycosis*, an invasive mycosis caused by *Talaromyces marneffei* (Tm), has rapidly increased in recent years, becoming an emerging pathogenic fungal disease. However, The driving factors and potential distribution of global *talaromycosis* is still unclear. Here, we developed maxent ecology model using environmental variables, Rhizomys distribution and HIV/AIDS epidemic to forecast ecological niche of *talaromycosis* worldwhile, as well as Identify the drivering factors. The constructed model had excellent performance with the area under the curve (AUC) of the receiver operating curve (ROC) of 0.997 in training data and 0.991 in testing data. Our model revealed that Rhizomys distribution, mean temperature of warmest quarter, precipitation of wettest month, HIV/AIDS epidemic and mean temperature of driest quarter were the top 5 important variables affecting *talaromycosis* distribution. In addition to traditional *talaromycosis* epidemic areas (South of the Yangtze River in China, Southeast Asian and North and Northeast India), our model also identified other potential epidemic regions, inculding parts of the North of the Yangtze River, Central America, West Coast of Africa, East Coast of South America, the Korean Peninsula and Japan. Our findings has redefined global *talaromycosis*, discovered hidden high-risk areas and prorvided insights about driving factors of *talaromycosis* distribution, which will help inform surveillance strategies and improve the effectiveness of public health interventions against Tm infections.

**Author Summary:** Our study aims to explore the spatial ecology of *talaromycosis* worldwhile. The diseases burden of *Talaromycosis*, a neglected zoonotic disease, is continuously rising in recent years because of the sheer size of susceptible population in the setting of increased globalization, rising HIV prevalence, and emerging iatrogenic immunodeficiency conditions. Here, we used historic reported *talaromycosis* cases from 1964 to 2017, combined with environmental factors, Rhizomys distribution and HIV/AIDS epidemic to build an maxent ecology model to define the ecological niche of *talaromycosis*, then predicting the potential distribution of the disease. The ecological niche of *talaromycosis* is characterized by a concentrated distribution, which can be cognitively divided into two regions: traditional talaromycosis epidemic areas (South of the Yangtze River in China, Southeast Asian and North and Northeast India), while other potential epidemic regions were predicted in parts of the North of the Yangtze River, Central America, West Coast of Africa, East Coast of South America, the Korean Peninsula and Japan. Our model also identified 5 driving factors affecting *talaromycosis* distribution. These findings will help demonstrate the global distribution of talaromycosis, discover hidden high-risk areas, and improve the effectiveness of public health interventions against Tm infections.

## Introduction

*Talaromyces marneffei* (Tm), originally known as *penicillium marneffei*, can cause a life-threatening fungal disease, *talaromycosis*. Most *talaromycosis* occur in patients with CD4^+^T cells count less than 100 cells/μL, especially HIV/AIDS patients(1, 2). Patients with *talaromycosis* usually suffer from weight loss, fever, hepatosplenomegaly, respiratory and gastrointestinal abnormalities(1–3). Rhizomys is the only known non-human host of Tm, and the Tm carried by the Rhizomys is genetically highly homologous to the strain in the patient(4–6). At present, although the reservoir of Tm in nature has not been fully clarified,, there is already evidence that occupational exposure to Rhizomys and its related soil is related to human infections(7).

*Talaromycosis* is a regional high-incidence opportunistic infection that is endemic throughout Southern China, Southeast Asia, and Northeastern India. The number of *talaromycosis* cases has rapidly increased due to the HIV epidemic, ranking 3^rd^ as the most common HIV-associated opportunistic infections and accounting for up to 16% of HIV admissions and is a leading cause of death in patients with advanced HIV disease in Thailand, Vietnam, and southern China(3, 6, 8). Due to lack of high-sensitivity early diagnosis technology, Tm infection leads to a high mortality. The mortality in patients with a disseminated disease without treatment reaches 80%-100%, and even with antifungal treatment, the mortality is as high as 30%(1, 2, 9, 10). Thus, due to intensified globalization, rising HIV/AIDS epidemic, and the emergence of iatrogenic immunodeficiency diseases, the number of *talaromycosis* is increasing year by year and continues to be a major public health problem.

*Talaromycosis* was not included in the disease surveillance system due to insufficient attention. Therefore, the epidemic regions of the disease can only be judged by the published primary cases. The traditional epidemic area of *talaromycosis* is generally considered to be in southern China, Southeast Asia and India(1, 2, 9, 10). A large number of studies have shown that there are primary cases of *talaromycosis* in Southeast Asian and northeastern India. However, there is no unified conclusion on the primary epidemic area in China(8), in addition to southern China, many cases have been reported in other regions of China. Thus, it is still unclear which regions have a higher risk of Tm transmission.

Also, what factors have contributed to the global distribution of *talaromycosis* has not been fully clarified. Previous studies have shown that the HIV/AIDS epidemic and the Rhizomys distribution are the driving factors affecting the distribution of *talaromycosis*(1, 6). Moreover, Tm can also be separated from Rhizomys feces and soil samples in its holes(11, 12). In addition, studies have shown that temperature and humidity are also related to the *talaromycosis* epidemic(13, 14). Therefore, related environmental factors may also be related to *talaromycosis* distribution.

To accurately predict *talaromycosis* distribution and analyze the driving factors, a large amount of data and high-performance ecological predictive model are required. In the field of ecology, niche models are often used to forecast and analyze the driving factors that affect the distribution of species and the best model currently is the Maximum Entropy (Maxent) model(13, 14). Maxent model is a species distribution model (SDM), originated from statistical mechanics(17). The Maxent model is easy to operate, the graphical interface is simple and clear, and the parameters are automatically set, which are favored by researchers(18, 19). This model has many successful applications in infectious diseases distribution prediction(20, 21).

In this study, we use the Maxent model to analyze the driving factors that affect *talaromycosis* distribution and predict the potential distribution of *talaromycosis* worldwide. This information is essential for making informed decisions around high-risk areas to implement targeted measures.

## Methods

### Rhizomys distribution information

Rhizomys can be broke down three different types of subclass: Rhizomys pruinosus Blyth, Rhizomys sinensis and Rhizomys sumatrensis. The distribution information of three kinds of Rhizomys were collected from Global Biodiversity Information Facility (GBIF) (Website: http://www.gbif.org/), which is one of the global specimen databases. 37,393 datasets with nearly 874,000,000 occurrence records from 1,135 publishing institutions have been collected in the database. In general, the occurrence records of Rhizomys from this database are relatively complete.

### *Talaromycosis* presence data

Point prevalence data for *talaromycosis* were collected from *talaromycosis* database constructed by our team. In brief, we collected the data of all laboratory-confirmed, non-duplicating *talaromycosis* published in the English and Chinese literature from the first case in 1964 to December 2017. The coordinate information of each case (only to the county level) is used for the further analysis.

### Worldwide HIV/AIDS epidemic

The HIV/AIDS epidemic came from the research of GBD 2015 HIV Collaborators, which containing the data of people living with HIV/AIDS from 1955 to 2015 in the world.

### Environmental variables information

The bioclimatic variables (bio1-bio19, see table 1 for details) were collected from WORLDCLIM databases, which is a free global climate data for ecological modeling and GIS. Bioclimatic variables were calculated from values of monthly rainfall and temperature, then generate new variables containing more biologically meaningful information. It is an authoritative dataset that often used in ecological modeling. Moreover, the worldwide leaf and tepography information came from Socioeconomic Data and Applications Center (SEDAC) databases (Website: http://sedac.ciesin.columbia.edu/), which is a data center in NASA’s Earth Observing System Data and Information System (EOSDIS) hosted by The Center for International Earth Science Information Network (CIESIN) at Columbia University.

**Table 1.**
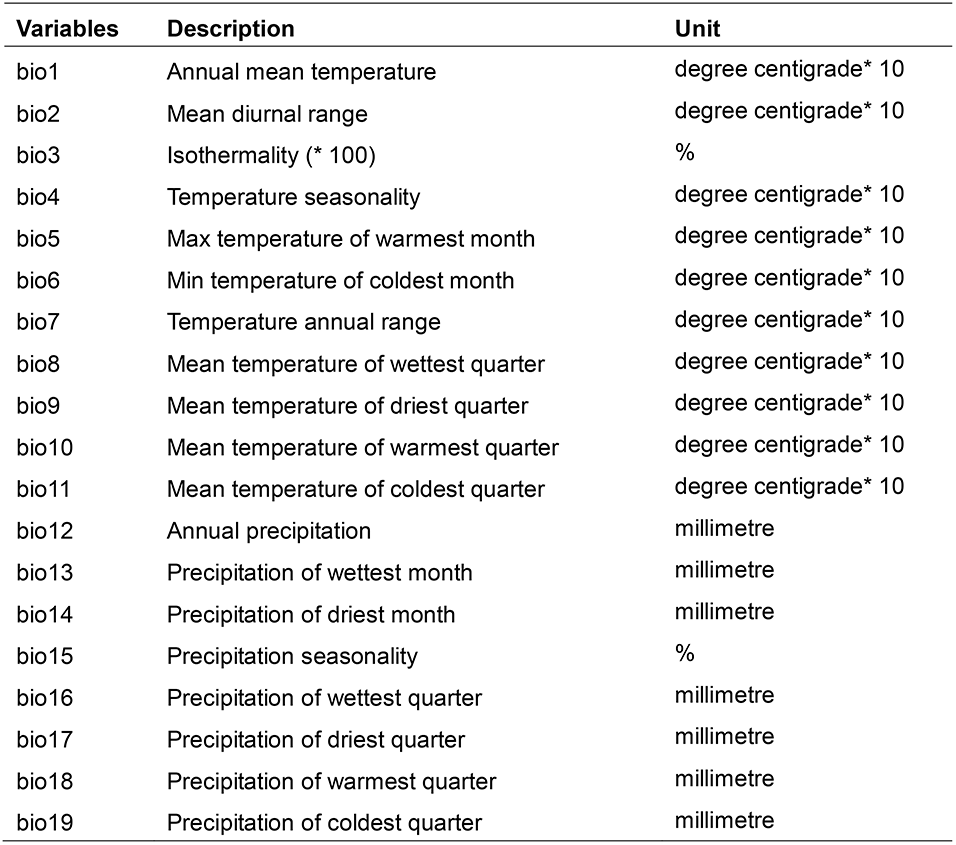
The bioclimatic variables used in construction of Maxent model

### Appropriate variables selection

Several highly correlation variables were used as the training part sets to develop the model, which may disturb the results of variables importance analysis, even the model performance. Thus, we use the multi-collinearity test (by Pearson Correlation Coefficient) to analyse the cross-correlation. If the Pearson Correlation Coefficient of two variables > 0.8 or −0.8, which meant they are highly correlation and one of them should be excluded. The one with high important gain should be retained, as it containing more helpful information to model contribution (Table 2).

**Table 2.** The bioclimatic variables used in construction of Maxent model for TM

| factor | bio1 | bio2 | bio3 | bio4 | bio5 | bio6 | bio7 | bio8 | bio9 | bio10 | bio11 | bio12 | bio13 | bio14 | bio15 | bio16 | bio17 | bio18 | bio19 | leaf | tepo | HIV | Rhizo |
| --- | --- | --- | --- | --- | --- | --- | --- | --- | --- | --- | --- | --- | --- | --- | --- | --- | --- | --- | --- | --- | --- | --- | --- |
| **bio1** | 1.00 |  |  |  |  |  |  |  |  |  |  |  |  |  |  |  |  |  |  |  |  |  |  |
| **bio2** | -0.33 | 1.00 |  |  |  |  |  |  |  |  |  |  |  |  |  |  |  |  |  |  |  |  |  |
| **bio3** | 0.74** | 0.08** | 1.00 |  |  |  |  |  |  |  |  |  |  |  |  |  |  |  |  |  |  |  |  |
| **bio4** | -0.93** | 0.24 | -0.87** | 1.00 |  |  |  |  |  |  |  |  |  |  |  |  |  |  |  |  |  |  |  |
| **bio5** | 0.60** | -0.14 | 0.15 | -0.29 | 1.00 |  |  |  |  |  |  |  |  |  |  |  |  |  |  |  |  |  |  |
| **bio6** | 0.98** | -0.43** | 0.76 | -0.96** | 0.47** | 1.00 |  |  |  |  |  |  |  |  |  |  |  |  |  |  |  |  |  |
| **bio7** | -0.92** | 0.43** | -0.79 | 0.98** | -0.26 | -0.98** | 1.00 |  |  |  |  |  |  |  |  |  |  |  |  |  |  |  |  |
| **bio8** | 0.68** | -0.30 | 0.31 | -0.46 | 0.68** | 0.61** | -0.49 | 1.00 |  |  |  |  |  |  |  |  |  |  |  |  |  |  |  |
| **bio9** | 0.98** | -0.29 | 0.74 | -0.92** | 0.58** | 0.97** | -0.91** | 0.66** | 1.00 |  |  |  |  |  |  |  |  |  |  |  |  |  |  |
| **bio10** | 0.72** | -0.41** | 0.20 | -0.42 | 0.93** | 0.63** | -0.46** | 0.80** | 0.69** | 1.00 |  |  |  |  |  |  |  |  |  |  |  |  |  |
| **bio11** | 0.99** | -0.31 | 0.81 | -0.98** | 0.47** | 0.99** | -0.96** | 0.59** | 0.97** | 0.60** | 1.00 |  |  |  |  |  |  |  |  |  |  |  |  |
| **bio12** | 0.71** | -0.49** | 0.50 | -0.70** | 0.30 | 0.74** | -0.73** | 0.43** | 0.68** | 0.45** | 0.72** | 1.00 |  |  |  |  |  |  |  |  |  |  |  |
| **bio13** | 0.67 | -0.33 | 0.49 | -0.66** | 0.31 | 0.67** | -0.65** | 0.48** | 0.64** | 0.42* | 0.68** | 0.88** | 1.00 |  |  |  |  |  |  |  |  |  |  |
| **bio14** | 0.26 | -0.37* | 0.25 | -0.25 | 0.08 | 0.32 | -0.32 | 0.13 | 0.28 | 0.21 | 0.27 | 0.50** | 0.09 | 1.00 |  |  |  |  |  |  |  |  |  |
| **bio15** | -0.12 | 0.51** | -0.03 | 0.13 | 0.03 | -0.21 | 0.24 | 0.09 | -0.11 | -0.09 | -0.14 | -0.31** | 0.13 | -0.74** | 1.00 |  |  |  |  |  |  |  |  |
| **bio16** | 0.66** | -0.33 | 0.46 | -0.66** | 0.31 | 0.66** | -0.65** | 0.43 | 0.63** | 0.41* | 0.67 | 0.90 | 0.99** | 0.11 | 0.08 | 1.00 |  |  |  |  |  |  |  |
| **bio17** | 0.31 | -0.39* | 0.29 | -0.29 | 0.13 | 0.37* | -0.37* | 0.17 | 0.33 | 0.26 | 0.32 | 0.53** | 0.14 | 0.99** | -0.73** | 0.15 | 1.00 |  |  |  |  |  |  |
| **bio18** | 0.15 | -0.40 | -0.13 | -0.14 | -0.15 | 0.18 | -0.23 | 0.25 | 0.12 | 0.08 | 0.15 | 0.48** | 0.42** | 0.22 | -0.03 | 0.44** | 0.21 | 1.00 |  |  |  |  |  |
| **bio19** | 0.46** | -0.22 | 0.61 | -0.49** | 0.15 | 0.51** | -0.52** | 0.25 | 0.47 | 0.24 | 0.48 | 0.51** | 0.27 | 0.79** | -0.50** | 0.24 | 0.81** | 0.00 | 1.00 |  |  |  |  |
| **leaf** | -0.05 | -0.33 | -0.13 | 0.03 | -0.10 | 0.03 | -0.05 | -0.08 | -0.03 | 0.00 | -0.03 | 0.02** | -0.13 | 0.35 | -0.27 | -0.14 | 0.32 | 0.13 | 0.10 | 1.00 |  |  |  |
| **tepo** | 0.12 | 0.15 | 0.49 | -0.30** | -0.30 | 0.18 | -0.27 | -0.21 | 0.15 | -0.28 | 0.20 | -0.10 | -0.15 | 0.17 | -0.04 | -0.15 | 0.17 | -0.23 | 0.20 | 0.21 | 1.00 |  |  |
| **HIV** | -0.07 | 0.62** | 0.04 | 0.03 | -0.04 | -0.13 | 0.14 | -0.14 | -0.02 | -0.18 | -0.06 | -0.32 | -0.30 | -0.21 | 0.16 | -0.26 | -0.23 | -0.18 | -0.20 | -0.17 | 0.09 | 1.00 |  |
| **Rhizo** | 0.15 | -0.45** | -0.15 | -0.17 | -0.21 | 0.20 | -0.28 | 0.07 | 0.13 | 0.00 | 0.16 | 0.29 | 0.26 | 0.00 | -0.03 | 0.31 | 0.00 | 0.81** | -0.13 | 0.14 | -0.03 | -0.14 | 1.00 |

Note: Correlations were significant at $\alpha = 0.05$ (\*\*p < 0.01, \*p < 0.05).

### Maxent model for *talaromycosis* Potential prevalence construction

To predict the potential distribution of *talaromycosis*, we used the maxent model (Website: http://www.cs.princeton.edu/~schapire/maxent), a predictive species potential distribution model based on niche principle. The models were constructed through the Software and running 10 times each model construction. 70% of the samples were randomly used as the training part to develop the model, while 30% of the points were selected for model testing. In order to get alternate estimates of variable importance, the jackknife analysis was employed. A series of models were developed, for each model run, each variable was excluded in turn, then a model created with the remaining variables. Furthermore, model was developed using each element in isolation. Variable with the highest gain when used in isolation appears to have the most useful information by itself. Variable decreases gain the most when it is omitted, therefore appearing to have the most information that is not present in other variables.

### Model validation

In the current study, two indicators (The receiver operating characteristic (ROC) curve was plotted and the area under the curve (AUC)) was calculated to show the sensitivity and specificity of model fitting and forecasting. AUC is usually divided into five categories, 0.5–0.6 in sufficient performance; 0.6–0.7 in poor performance; 0.7–0.8 in average performance; 0.8–0.9 in good performance; and 0.9–1.0 in excellent performance(22).

## Result

### Rhizomys prediction map

The environmental variables (n=21) were first used to develop a Rhizomys prediction model, jackknife algorithm was used to calculate the importance of each variable. Combine the results of the Spearman correlation matrix, the variables (bio2, bio3, bio5, bio7, bio8, bio9, bio15, bio18, bio19, leaf and tepograph) that can be included in the model were filter out (Appendix 1, SFig. 1 and STable 1). Then use selected variables to build a maxent model to predict the distribution of Rhizomys (Appendix 1, SFig. 2).

The results show that South China, Southeast Asia, Central America, West Coast of Africa, North India, East Coast of India, East Coast of Australia, Korean Peninsula and Japan are potential distribution areas for Rhizomys (Appendix 1, SFig. 3).

### Importance of variables for *talaromycosis* distribution prediction

Similar to the Rhizomys distribution prediction, all variables (n=23) were used to construct a maxent model and calculate the importance of each variable. And select the appropriate independent variables based on the Spearman correlation matrix. Finally, bio2, bio3, bio9, bio10, bio13, bio15, leaf, tepograph, HIV/AIDS epidemic and Rhizomys distribution were used to the further modeling (Figure 1).

**Fig. 1.**
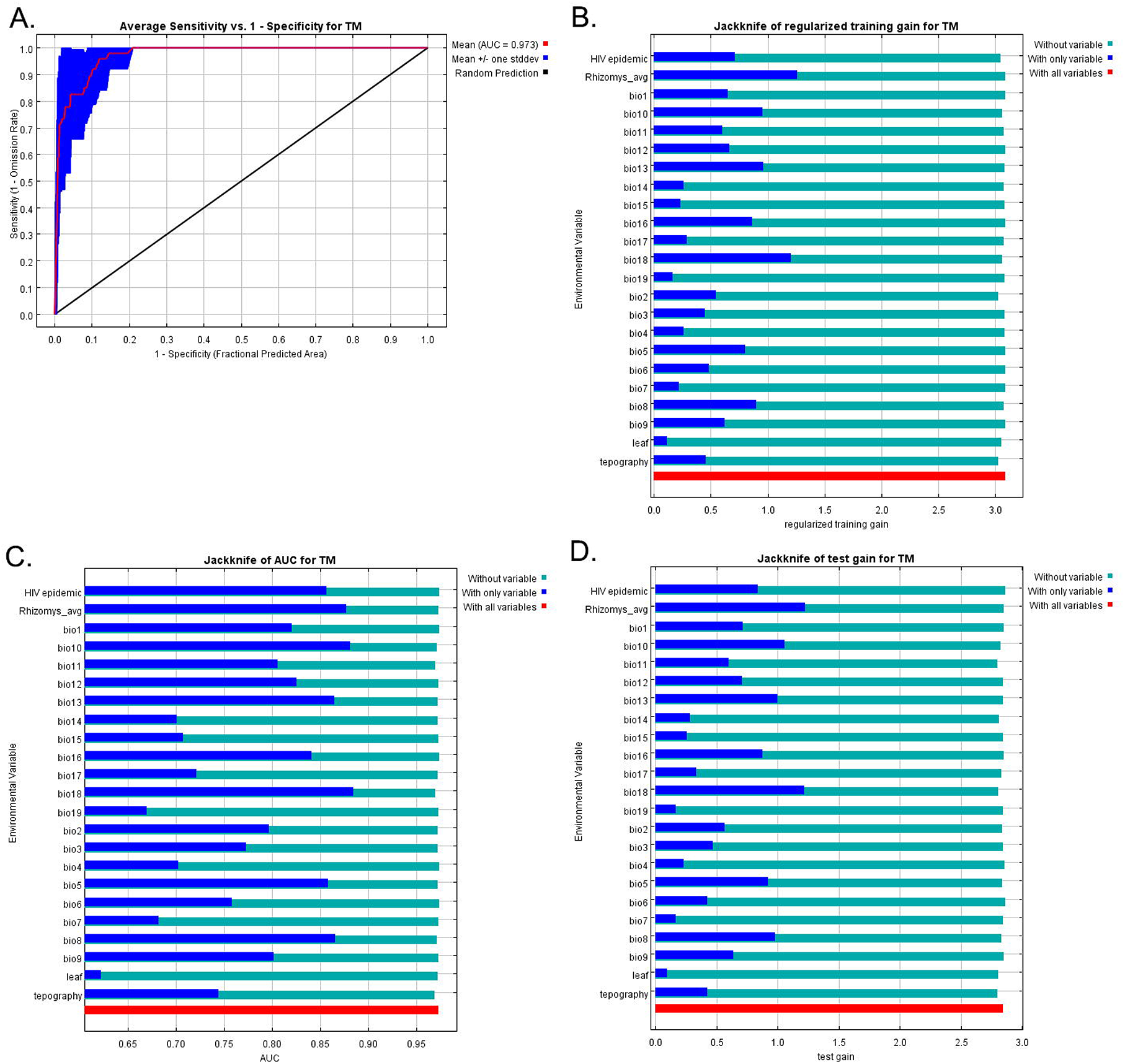
The performance of first maxent model for *talaromycosis*. (A.) The ROC curve of maxent model. (B.) Jackknife analysis of regularized training gain for 10 runs. (C.) Jackknife analysis of test gain for 10 runs. (D.) Jackknife analysis of regularized training gain for 10 runs.

### Maxent model of *talaromycosis*

Figure 2 showed the potential distribution of *talaromycosis* during the study period. The resulting maps show that Southern China, Southeast Asia, Central America, West coast of Africa, northern India, eastern coast of India, eastern coast of Australia, the Korean peninsula, and Japan were the potential prevalence regions of *talaromycosis.* Figure 3A showed that the AUC values of training data and testing data were 0.987 and 0.991, respectively, which indicated that the simulation was excellent and proved that the model can be used to predict the potential distribution of *talaromycosis* in the world.

**Fig. 2.**
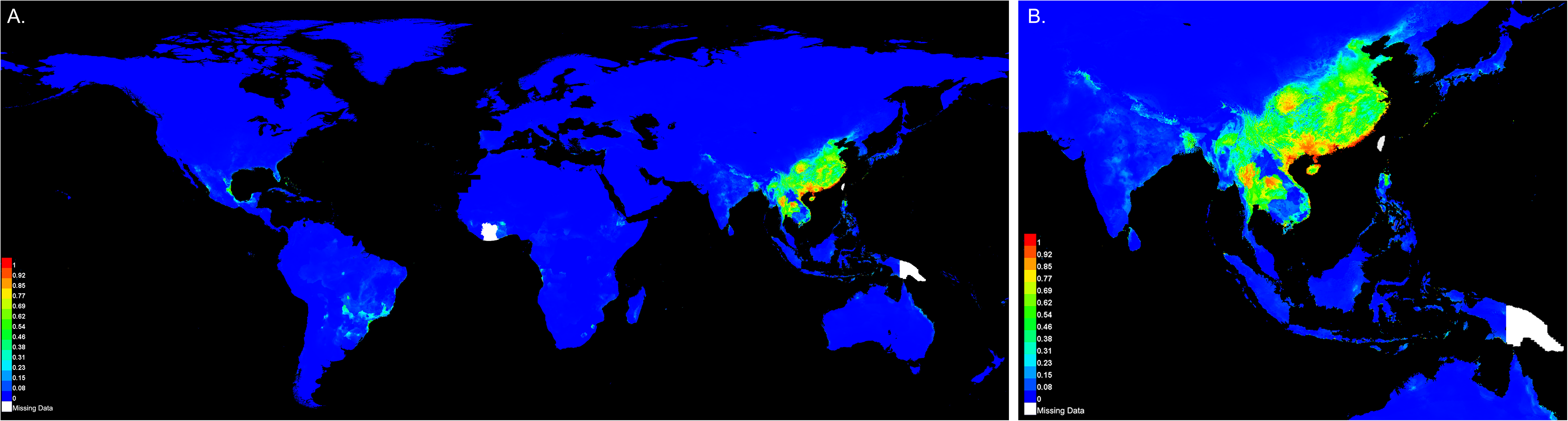
The potential distribution map of *talaromycosis*.

**Fig. 3.**
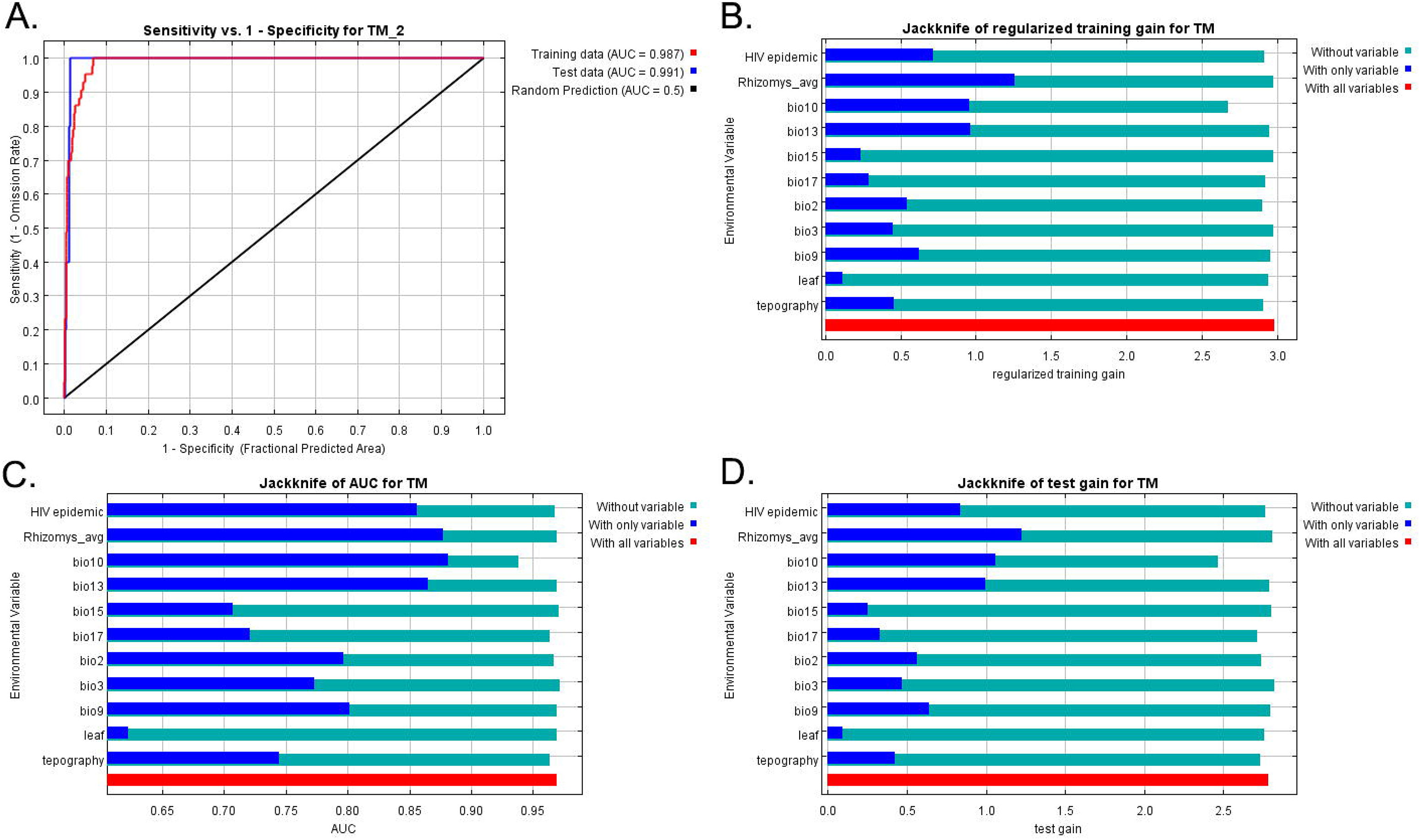
The performance of final maxent model for *talaromycosis*. (A.) The ROC curve of maxent model. (B.) Jackknife analysis of regularized training gain for 10 runs. (C.) Jackknife analysis of test gain for 10 runs. (D.) Jackknife analysis of regularized training gain for 10 runs.

### Contribution of the variables to models

Jackknife test showed that Rhizomys distribution, mean temperature of warmest quarter, precipitation of wettest month, HIV/AIDS epidemic and mean temperature of driest quarter were the top 5 important variables for *talaromycosis* potential prevalence, which the regularized training and test gain were significantly bigger than 0.5 (Figure 3B-D). And the rest of variables were less importance to the model development.

### Variables affecting the occurrence of *talaromycosis*

Mean temperature of warmest quarter, precipitation of wettest month, HIV/AIDS epidemic and mean temperature of driest quarter showed similar patterns, in which the probability of *talaromycosis* presence was initially increased, then reached a peak, and subsequently decreased. For mean temperature of warmest quarter, probability of *talaromycosis* presence increased from 15°C, peaking at 28°C and then decreased to 35°C. For precipitation of wettest month, the probability increased from 0mm, peaking at 200mm and then decreased to 1200mm. For HIV/AIDS epidemic, the probability increased from 500,000, peaking at 800,000 and then decreased to 5,200,000. For mean temperature of driest quarter, the probability increased from −20°C, peaking at 8°C and then decreased to 35°C. Probability of Rhizomys occurrence was positively associated with *talaromycosis* presence (Figure 4).

**Fig. 4.**
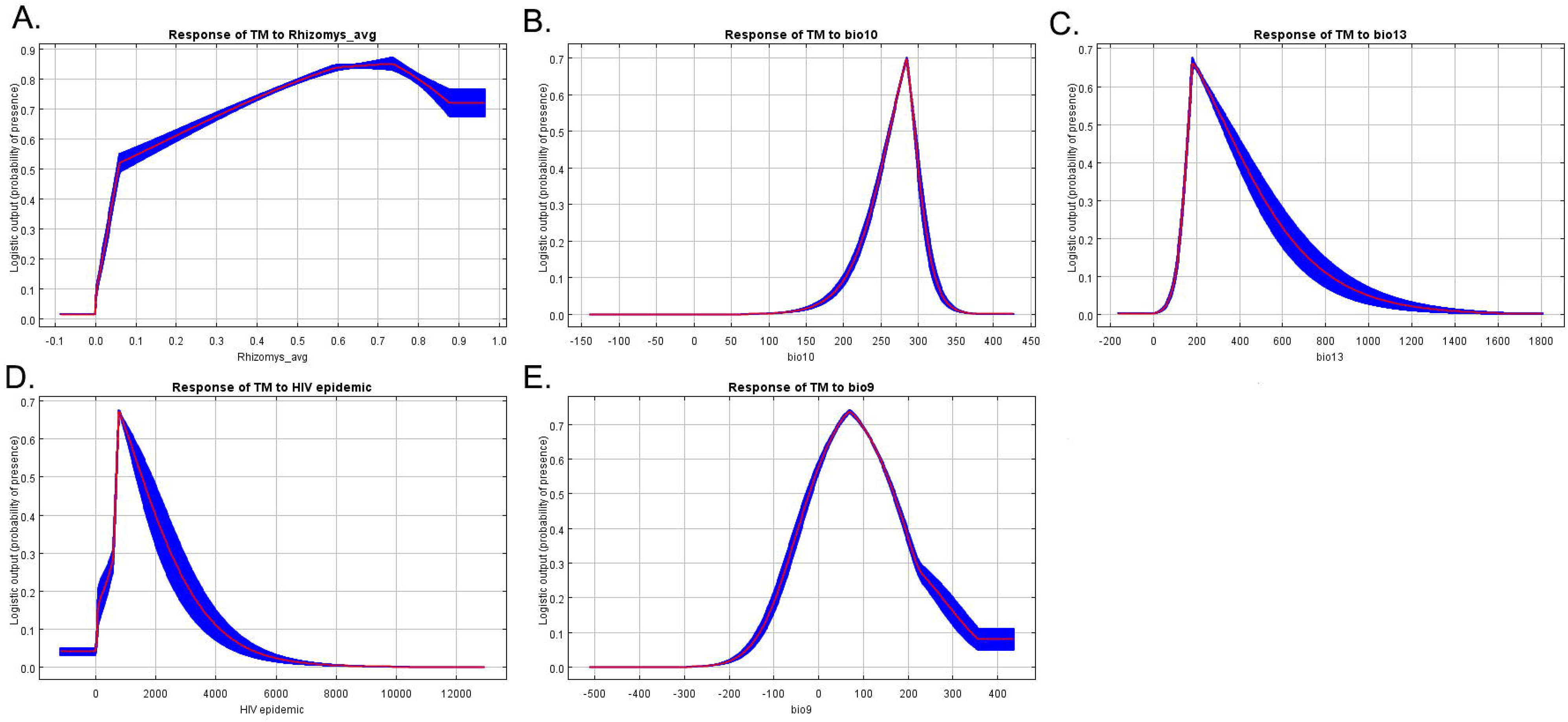
Response curves of variables associated with *talaromycosis*. The curves show the mean response of the 10 replicate Maxent runs (red) and the mean +/− one standard deviation (blue, two shades for categorical variables).

## Discussion

In this study, we predicted the potential global *talaromycosis* distribution. The results showed that *talaromycosis* potential epidemic regions are mainly distributed in South China, Southeast Asia, Central America, the west coast of Africa, northern India, the east coast of India, the east coast of Australia, the Korean Peninsula and Japan. Furthermore, we found that the distribution of Rhizomys, the average temperature of the warmest season, the precipitation of the wettest month, the average temperature of the driest season and the HIV/AIDS prevalence are the top five important driving factors for the potential prevalence of *talaromycosis*.

As a zoonotic fungal disease, the ecological distribution of the animal host largely determines the epidemiological distribution of the disease. Rhizomys is the only known non-human host of Tm, which the frequency of asymptomatic infection is very high(4, 5). Although there is no direct evidence that Tm is directly transmitted to humans through Rhizomys, studies have shown that Tm isolates from Rhizomys and humans have similar or identical genotypes(4, 5). Our findings also provide ecological evidence for this association. The results of the Maxent model show that the distribution of Rhizomys is the most important driving factor for the distribution of *talaromycosis.* Rhizomys, commonly known as bamboo rat, can be divided into three subspecies, Rhizomys pruinosus, Rhizomys sinensis and Rhizomys sumatrensis, which mainly inhabit tropical and subtropical forests, shrubs and bamboo forests in Southern China, Southern Asia, and Africa. This may partly explain that China and Southeast Asia are currently the main endemic areas of *talaromycosis*. Interestingly, even if Rhizomys are also distributed in Africa, it is not considered that Africa is also the primary endemic area of *talaromycosis*. At present, there is no relevant research on the Tm carrying rate in African Rhizomys, which is very necessary for further identification of *talaromycosis* potential endemic areas.

The first and second naturally occurring human case of *talaromycosis* was reported in 1973 and 1984, respectively(23, 24). Although there were several reports of cases after a period of time, the epidemic did not seem to be very serious(25). With the global HIV/AIDS pandemic reached Rhizomys habitats such as China, Southeast Asia, and India, a large number of human *talaromycosis* were reported(26–32). *Talaromycosis* has been regarded as an important opportunistic infection of HIV/AIDS patients, which key mechanism is the exhaustion of host CD4+ T cells caused by HIV infection(33–36). Without CD4+ T cells, the host’s innate immune cells, such as macrophages and neutrophils, are difficult to inhibit Tm infection(37–42). More importantly, it is difficult to form or maintain granulomas at the infected site, which can easily cause the spread of Tm infection(43). Our model results showed that HIV/AIDS epidemic was also a key driving factor that effecting the *talaromycosis* distribution. However, the response curve shows that it is not that the more serious the HIV/AIDS epidemic, the higher the probability of *talaromycosis*. HIV infection only caused an increase in the local susceptible population. Thus, in some areas where Rhizomys are lacking, such as southern Africa, even if the HIV/AIDS epidemic is very serious, there are currently no case reports.

*Talaromycosis* occurs in a tropical or subtropical monsoon climate with unique hydrological characteristics, which is very suitable for the survival of Tm natural host Rhizomys(6). Therefore, environmental factors are also important driving factors affecting the distribution of *talaromycosis*. The association has been discovered in a previous study from Vietnam, which showed that *talaromycosis* incidence increases 30–50% during the rainy months(1, 14), and also correlation with humidity level(2, 13). We also got similar research results through the niche model, which showed that temperature and precipitation can be used as the predicted factors for *talaromycosis* distribution. First, precipitation and temperature may directly affect the niche of Rhizomys. The ecological model results showed that precipitation and temperature are both important driving factors affecting the distribution of Rhizomys. Secondly, precipitation can promote the growth of Tm and fully expose the Tm spores in the soil, leading to the expansion of its natural reservoir. At the same time, the host’s exposure probability also increases. Unfortunately, due to the lack of humidity variable in environmental variable data, we have not analyzed how much humidity contributes to the distribution of *talaromycosis*. Since its natural reservoir and transmission mode are not yet clear, it needs to be further evaluated in epidemiological studies.

It is worth noting that Lo, Patassi and Guiguemde respectively reported *talaromycosis* patients in Ghana, Togo and Ouagadougou (Burkina Faso), all of whom had never been to Asia and denied having contact with animals with lesions(44–46). These countries are located in western Africa and are neighbors to each other. The prediction results of the Maxent niche model show that these regions belong to the potential epidemic area of *talaromycosis*. Therefore, we speculate that western Africa are another region of *talaromycosis* outside of the traditional epidemic regions. In western Africa, Tm infection may be not known to clinicians. The insufficient technology platform for mycological diagnosis may lead to missed diagnosis of local *talaromycosis.* Therefore, medical staff in areas with severe HIV/AIDS epidemics should pay more attention to Tm infection. In addition, it is urgent to establish an efficient and sensitive rapid diagnosis technology for *talaromycosis*.

In this study, the limitations of our analysis should be acknowledged. First, the data comes from published literature, which will lead to publication bias; second, the accuracy of the geographic coordinates of *talaromycosis* we obtained is limited, resulting in insufficient sample size; third, the model does not consider other causes of autoimmune suppression, such as the use of immunosuppressive agents, IFN-γ gene mutations, etc. Unfortunately, we are unable to obtain complete and up-to-date data on these variables, which should be considered in future work.

## Conclusion

This study produced the potential global distribution of *talaromycosis* and analyzes the driving factors affecting its presence. The findings will help to better understand the ecology of global *talaromycosis*, remind people with low immunity in potential epidemic areas to pay attention to Tm infection, and may develop effective *talaromycosis* prevention and control strategies for the epidemic regions.

## Acknowledgment

We thank the researchers who provided free niche data online. The study was supported by National Natural Science Foundation of China (NSFC; 81971934, 81760602), Thousands of Young and Middle-aged Key Teachers Training Program in Guangxi Colleges and Universities (to Junjun Jiang), Guangxi Medical University Training Program for Distinguished Young Scholars (to Junjun Jiang), Guangxi Medical University Youth Science Fund Project (GXMUYSF202106, to Wudi Wei) and China Postdoctoral Science Foundation (2020M683212, to Wudi Wei).

## Author Contributions

LH and YL designed the research. WW, WG and LJ collected and cleaned the data. WW and CN analyzed the data. WW, CN, and XB constructed the mathematical model. JJ visualized the results. WW, CN and JH wrote the initial draft manuscript. All authors have seen, edited, and approved the final version of the manuscript.

## Supporting Information

**Appendix 1. Niche model construction for Rhizomys prediction map.**

## References

1. Le T, Wolbers M, Chi NH, Quang VM, Chinh NT, Lan NP, et al. Epidemiology, seasonality, and predictors of outcome of AIDS-associated Penicillium marneffei infection in Ho Chi Minh City, Viet Nam. Clin Infect Dis. 2011;52(7):945–52. http://doi.org/10.1093/cid/cir028

2. Larsson M, Nguyen LH, Wertheim HF, Dao TT, Taylor W, Horby P, et al. Clinical characteristics and outcome of Penicillium marneffei infection among HIV-infected patients in northern Vietnam. Aids Res Ther. 2012;9(1):24. http://doi.org/10.1186/1742-6405-9-24

3. Supparatpinyo K, Khamwan C, Baosoung V, Nelson KE, Sirisanthana T. Disseminated Penicillium marneffei infection in southeast Asia. Lancet (London, England). 1994;344(8915):110–3. http://doi.org/10.1016/s0140-6736(94)91287-4

4. Gugnani H, Fisher MC, Paliwal-Johsi A, Vanittanakom N, Singh I, Yadav PS. Role of Cannomys badius as a natural animal host of Penicillium marneffei in India. J Clin Microbiol. 2004;42(11):5070–5. http://doi.org/10.1128/JCM.42.11.5070-5075.2004

5. Cao C, Liang L, Wang W, Luo H, Huang S, Liu D, et al. Common reservoirs for Penicillium marneffei infection in humans and rodents, China. Emerg Infect Dis. 2011;17(2):209–14. http://doi.org/10.3201/eid1702.100718

6. Limper AH, Adenis A, Le T, Harrison TS. Fungal infections in HIV/AIDS. The Lancet. Infectious diseases. 2017;17(11):e334–43. http://doi.org/10.1016/S1473-3099(17)30303-1

7. Chariyalertsak S, Sirisanthana T, Supparatpinyo K, Praparattanapan J, Nelson KE. Case-control study of risk factors for Penicillium marneffei infection in human immunodeficiency virus-infected patients in northern Thailand. Clin Infect Dis. 1997;24(6):1080–6. http://doi.org/10.1086/513649

8. Vanittanakom N, Cooper CJ, Fisher MC, Sirisanthana T. Penicillium marneffei infection and recent advances in the epidemiology and molecular biology aspects. Clin Microbiol Rev. 2006;19(1):95–110. http://doi.org/10.1128/CMR.19.1.95-110.2006

9. Sirisanthana T, Supparatpinyo K. Epidemiology and management of penicilliosis in human immunodeficiency virus-infected patients. Int J Infect Dis. 1998;3(1):48–53. http://doi.org/10.1016/s1201-9712(98)90095-9

10. Jiang J, Meng S, Huang S, Ruan Y, Lu X, Li JZ, et al. Effects of Talaromyces marneffei infection on mortality of HIV/AIDS patients in southern China: a retrospective cohort study. Clin Microbiol Infect. 2019;25(2):233–41. http://doi.org/10.1016/j.cmi.2018.04.018

11. Chariyalertsak S, Vanittanakom P, Nelson KE, Sirisanthana T, Vanittanakom N. Rhizomys sumatrensis and Cannomys badius, new natural animal hosts of Penicillium marneffei. J Med Vet Mycol. 1996;34(2):105–10.

12. Huang X, He G, Lu S, Liang Y, Xi L. Role of Rhizomys pruinosus as a natural animal host of Penicillium marneffei in Guangdong, China. Microb Biotechnol. 2015;8(4):659–64. http://doi.org/10.1111/1751-7915.12275

13. Bulterys PL, Le T, Quang VM, Nelson KE, Lloyd-Smith JO. Environmental predictors and incubation period of AIDS-associated penicillium marneffei infection in Ho Chi Minh City, Vietnam. Clinical infectious diseases : an official publication of the Infectious Diseases Society of America. 2013;56(9):1273–9. http://doi.org/10.1093/cid/cit058

14. Chariyalertsak S, Sirisanthana T, Supparatpinyo K, Nelson KE. Seasonal variation of disseminated Penicillium marneffei infections in northern Thailand: a clue to the reservoir? J Infect Dis. 1996;173(6):1490–3. http://doi.org/10.1093/infdis/173.6.1490

15. Sofizadeh A, Rassi Y, Vatandoost H, Hanafi-Bojd AA, Mollalo A, Rafizadeh S, et al. Predicting the Distribution of Phlebotomus papatasi (Diptera: Psychodidae), the Primary Vector of Zoonotic Cutaneous Leishmaniasis, in Golestan Province of Iran Using Ecological Niche Modeling: Comparison of MaxEnt and GARP Models. J Med Entomol. 2017;54(2):312–20. http://doi.org/10.1093/jme/tjw178

16. Srivastava V, Griess VC, Keena MA. Assessing the Potential Distribution of Asian Gypsy Moth in Canada: A Comparison of Two Methodological Approaches. Sci Rep-Uk. 2020;10(1):22. http://doi.org/10.1038/s41598-019-57020-7

17. Haegeman B, Etienne RS. Entropy maximization and the spatial distribution of species. The American naturalist. 2010;175(4):E74–90. http://doi.org/10.1086/650718

18. Wang XY, Huang XL, Jiang LY, Qiao GX. Predicting potential distribution of chestnut phylloxerid (Hemiptera: Phylloxeridae) based on GARP and Maxent ecological niche models. J Appl Entomol. 2010;134(1):45–54.

19. Narouei-Khandan HA, Halbert SE, Worner SP, van Bruggen AHC. Global climate suitability of citrus huanglongbing and its vector, the Asian citrus psyllid, using two correlative species distribution modeling approaches, with emphasis on the USA. Eur J Plant Pathol. 2016;144(3):655–70. http://doi.org/10.1007/s10658-015-0804-7

20. Wang L, Hu W, Soares Magalhaes RJ, Bi P, Ding F, Sun H, et al. The role of environmental factors in the spatial distribution of Japanese encephalitis in mainland China. Environ Int. 2014;73:1–9. http://doi.org/10.1016/j.envint.2014.07.004

21. Gao X, Cao Z. Meteorological conditions, elevation and land cover as predictors for the distribution analysis of visceral leishmaniasis in Sinkiang province, Mainland China. Sci Total Environ. 2019;646:1111–6. http://doi.org/10.1016/j.scitotenv.2018.07.391

22. Yunsheng W. Application of ROC curve analysis in evaluating the performance of alien species’ potential distribution models. Biodiversity Science. 2007;15. http://doi.org/10.1360/biodiv.060280

23. DiSalvo AF, Fickling AM, Ajello L. Infection caused by Penicillium marneffei: description of first natural infection in man. Am J Clin Pathol. 1973;60(2):259–63. http://doi.org/10.1093/ajcp/60.2.259

24. Pautler KB, Padhye AA, Ajello L. Imported penicilliosis marneffei in the United States: report of a second human infection. Sabouraudia. 1984;22(5):433–8. http://doi.org/10.1080/00362178485380691

25. Deng Z, Ribas JL, Gibson DW, Connor DH. Infections caused by Penicillium marneffei in China and Southeast Asia: review of eighteen published cases and report of four more Chinese cases. Rev Infect Dis. 1988;10(3):640–52. http://doi.org/10.1093/clinids/10.3.640

26. Liao X, Ran Y, Chen H, Meng W, Xiang B, Kang M, et al. [Disseminated Penicillium marneffei infection associated with AIDS, report of a case]. Zhonghua Yi Xue Za Zhi. 2002;82(5):325–9.

27. R. Bailloud MSBS. Premiers cas d’infection à Penicillium marneffei identifiés chez l’immunodéprimé au Cambodge. Journal de Mycologie Médicale. 2002;1046(3):101–51. http://dx.doi.org/JMM-09-2002-12-3-1156-5233-101019-ART8

28. Rokiah I, Ng KP, Soo-Hoo TS. Penicillium marneffei infection in an AIDS patient--a first case report from Malaysia. Med J Malaysia. 1995;50(1):101–4.

29. Hien TV, Loc PP, Hoa NT, Duong NM, Quang VM, McNeil MM, et al. First cases of disseminated penicilliosis marneffei infection among patients with acquired immunodeficiency syndrome in Vietnam. Clin Infect Dis. 2001;32(4):e78–80. http://doi.org/10.1086/318703

30. Ranjana KH, Priyokumar K, Singh TJ, Gupta C, Sharmila L, Singh PN, et al. Disseminated Penicillium marneffei infection among HIV-infected patients in Manipur state, India. J Infect. 2002;45(4):268–71. http://doi.org/10.1053/jinf.2002.1062

31. Ko KF. Retropharyngeal abscess caused by Penicillium marneffei: an unusual cause of upper airway obstruction. Otolaryngol Head Neck Surg. 1994;110(4):445–6. http://doi.org/10.1177/019459989411000417

32. Wong KH, Lee SS, Chan KC, Choi T. Redefining AIDS: case exemplified by Penicillium marneffei infection in HIV-infected people in Hong Kong. Int J Std Aids. 1998;9(9):555–6.

33. Supparatpinyo K, Chiewchanvit S, Hirunsri P, Uthammachai C, Nelson KE, Sirisanthana T. Penicillium marneffei infection in patients infected with human immunodeficiency virus. Clin Infect Dis. 1992;14(4):871–4. http://doi.org/10.1093/clinids/14.4.871

34. Kudeken N, Kawakami K, Kusano N, Saito A. Cell-mediated immunity in host resistance against infection caused by Penicillium marneffei. J Med Vet Mycol. 1996;34(6):371–8. http://doi.org/10.1080/02681219680000671

35. Kudeken N, Kawakami K, Saito A. CD4+ T cell-mediated fatal hyperinflammatory reactions in mice infected with Penicillium marneffei. Clin Exp Immunol. 1997;107(3):468–73. http://doi.org/10.1046/j.1365-2249.1997.d01-945.x

36. Viviani M, Hill J, Dixon D. Penicillium Marneffei: Dimorphism and Treatment. 1993:{}. http://doi.org/10.1007/978-1-4615-2834-0_37

37. Cogliati M, Roverselli A, Boelaert JR, Taramelli D, Lombardi L, Viviani MA. Development of an in vitro macrophage system to assess Penicillium marneffei growth and susceptibility to nitric oxide. Infect Immun. 1997;65(1):279–84. http://doi.org/10.1128/IAI.65.1.279-284.1997

38. Kudeken N, Kawakami K, Saito A. Different susceptibilities of yeasts and conidia of Penicillium marneffei to nitric oxide (NO)-mediated fungicidal activity of murine macrophages. Clin Exp Immunol. 1998;112(2):287–93. http://doi.org/10.1046/j.1365-2249.1998.00565.x

39. Kudeken N, Kawakami K, Saito A. Role of superoxide anion in the fungicidal activity of murine peritoneal exudate macrophages against Penicillium marneffei. Microbiol Immunol. 1999;43(4):323–30. http://doi.org/10.1111/j.1348-0421.1999.tb02412.x

40. Kudeken N, Kawakami K, Saito A. Cytokine-induced fungicidal activity of human polymorphonuclear leukocytes against Penicillium marneffei. FEMS Immunol Med Microbiol. 1999;26(2):115–24. http://doi.org/10.1111/j.1574-695X.1999.tb01378.x

41. Levitz SM. Overview of host defenses in fungal infections. Clin Infect Dis. 1992;14 Suppl 1:S37–42. http://doi.org/10.1093/clinids/14.supplement_1.s37

42. Taramelli D, Brambilla S, Sala G, Bruccoleri A, Tognazioli C, Riviera-Uzielli L, et al. Effects of iron on extracellular and intracellular growth of Penicillium marneffei. Infect Immun. 2000;68(3):1724–6. http://doi.org/10.1128/iai.68.3.1724-1726.2000

43. Pagán AJ, Ramakrishnan L. The Formation and Function of Granulomas. Annu Rev Immunol. 2018;36:639–65. http://doi.org/10.1146/annurev-immunol-032712-100022

44. Lo Y, Tintelnot K, Lippert U, Hoppe T. Disseminated Penicillium marneffei infection in an African AIDS patient. Trans R Soc Trop Med Hyg. 2000;94(2):187. http://doi.org/10.1016/s0035-9203(00)90271-2

45. Guiguemde KT, Sawadogo PM, Zida A, Cisse M, Sangare I, Bamba S. First case report of Talaromyces marneffei infection in HIV-infected patient in the city of Ouagadougou (Burkina Faso). Medical Mycology Case Reports. 2019;26:10–2. http://doi.org/10.1016/j.mmcr.2019.09.003

46. Patassi AA, Saka B, Landoh DE, Kotosso A, Mawu K, Halatoko WA, et al. First observation in a non-endemic country (Togo) of Penicillium marneffei infection in a human immunodeficiency virus-infected patient: a case report. BMC Res Notes. 2013;6:506. http://doi.org/10.1186/1756-0500-6-506

